# The landscape of extrachromosomal circular DNA in medulloblastoma

**DOI:** 10.1101/2021.10.18.464907

**Authors:** Owen S Chapman, Jens Luebeck, Sameena Wani, Ashutosh Tiwari, Meghana Pagadala, Shanqing Wang, Jon D Larson, Joshua T Lange, Ivy Tsz-Lo Wong, Siavash R Dehkordi, Sahaana Chandran, Miriam Adam, Yingxi Lin, Edwin Juarez, James T Robinson, Sunita Sridhar, Denise M Malicki, Nicole Coufal, Michael Levy, John R Crawford, Scott L Pomeroy, Jeremy Rich, Richard H Scheuermann, Hannah Carter, Jesse Dixon, Paul S Mischel, Ernest Fraenkel, Robert J Wechsler-Reya, Vineet Bafna, Jill P Mesirov, Lukas Chavez

**Affiliations:** Bioinformatics and Systems Biology Graduate Program, University of California San Diego, San Diego, CA, USA; Department of Biomedical Informatics, University of California San Diego, San Diego, CA, USA; Department of Medicine, University of California San Diego, San Diego, CA, USA; Department of Computer Science and Engineering, University of California San Diego, San Diego, CA, USA; Sanford Burnham Prebys Medical Discovery Institute, San Diego, CA, USA; Medical Scientist Training Program, University of California San Diego, San Diego, CA, USA; Biomedical Science Graduate Program, University of California San Diego, San Diego, CA, USA; Department of Pathology, Stanford University School of Medicine, Stanford, CA, USA; ChEM-H, Stanford University, Stanford, CA, USA; Salk Institute for Biological Studies, La Jolla, CA, USA; Department of Biological Engineering, Massachusetts Institute of Technology, Cambridge, MA, USA; Department of Neurosciences and Pediatrics, UC San Diego and Rady Children’s Hospital, San Diego, CA, USA; Division of Pathology, UC San Diego and Rady Children’s Hospital, San Diego, CA, USA; Department of Neurology, Boston Children’s Hospital, Boston, MA, USA; Harvard Medical School, Boston, MA, USA; Eli and Edythe Broad Institute of MIT and Harvard, Cambridge, MA, USA; J. Craig Venter Institute, La Jolla, CA, USA; Halıcıoğlu Data Science Institute, University of California San Diego, San Diego, CA, USA; Moores Cancer Center, University of California San Diego, San Diego, CA, USA

## Abstract

Extrachromosomal circular DNA (ecDNA) is an important driver of aggressive tumor growth, promoting high oncogene copy number, intratumoral heterogeneity, accelerated evolution of drug resistance, enhancer rewiring, and poor outcome. ecDNA has been reported in medulloblastoma (MB), the most common malignant pediatric brain tumor, but the ecDNA landscape and its association with specific MB subgroups, its impact on enhancer rewiring, and its potential clinical implications, are not known. We assembled a retrospective cohort of 468 MB patient samples with available whole genome sequencing (WGS) data covering the four major MB subgroups WNT, SHH, Group 3 and Group 4. Using computational methods for the detection and reconstruction of ecDNA^1^, we find ecDNA in 82 patients (18%) and observe that ecDNA+ MB patients are more than twice as likely to relapse and three times as likely to die of disease. In addition, we find that individual medulloblastoma tumors often harbor multiple ecDNAs, each containing different amplified oncogenes along with co-amplified non-coding regulatory enhancers. ecDNA was substantially more prevalent among 31 analyzed patient-derived xenograft (PDX) models and cell lines than in our patient cohort. By mapping the accessible chromatin and 3D conformation landscapes of MB tumors that harbor ecDNA, we observe frequent candidate “enhancer rewiring” events that spatially link oncogenes with co-amplified enhancers. Our study reveals the frequency and diversity of ecDNA in a subset of highly aggressive tumors and suggests enhancer rewiring as a frequent oncogenic mechanism of ecDNAs in MB. Further, these results demonstrate that ecDNA is a frequent and potent driver of poor outcome in MB patients.

## INTRODUCTION

Extrachromosomal circular DNA molecules (ecDNA), also known as double minutes (dm), have been described in isolated tumor and tumor-derived cells since the 1960s^2^, but recent results have shown ecDNA to be far more common in human cancer than previously assumed^3,4^. Commonly defined as a circular, acentric chromatin body hundreds of kilobases to tens of megabases in length, ecDNA is now understood to have prominent roles in oncogenesis, tumor evolution, and chemotherapeutic resistance^5^. Algorithmic advances, and the increasing availability of whole genome sequencing (WGS) data from human tumors, have facilitated statistical analyses of ecDNA across a large cohort of adult tumors^4^. Studies in several cancers, including adult glioblastoma^6^, pediatric neuroblastoma^7^, acute myeloid leukemia^8^, and *HER2+* breast cancer^9^, have identified ecDNA as a prognostic biomarker for poor outcome. ecDNA has been shown to allow amplified oncogenes to “hijack” non-coding regulatory enhancers which would be inaccessible to them under normal karyotypic topology^10-12^. It has also been shown that ecDNA catalyzes rapid acquisition of drug resistance in response to chemotherapy^13-15^. However, the prevalence, diversity, and composition of ecDNA, including the role of non-coding regulatory DNA in transcriptional activation of co-amplified oncogenes, has not been studied in medulloblastoma (MB), the most common malignant pediatric brain tumor.

Medulloblastomas were represented among the first patient case reports describing ecDNA^2,16^, and several patient-derived cell line models of MB are known to contain ecDNA^12,17^. Few effective targeted molecular treatments exist for MB, and the current standard of care carries substantial risk of developmental disorder, neurological damage, and secondary metastasis^18^. There are four major molecular subgroups of MB: WNT, SHH, Group 3 (G3) and Group 4 (G4)^19^. Prognosis is especially poor for a subset of aggressive MYC-activated Group 3 tumors, and for p53-mutant SHH subgroup tumors^20,21^. Both *MYC* amplification and p53 mutation have been co-reported alongside ecDNA in MB^22,23^, but the strength of any possible association has not yet been quantified in a patient population. Although the genomic landscape of medulloblastoma subgroups has been largely characterized^20^, it remains unknown how frequently ecDNA occurs in MB patients, and whether the presence of ecDNA may affect prognosis or correlate with other molecular features.

## RESULTS

### ecDNA in medulloblastoma patients

To examine the landscape of ecDNA in medulloblastoma, we accessed WGS data available in three cancer cloud genomics platforms: St Jude Cloud^24^, the Childrens’ Brain Tumor Network (CBTN), and the International Cancer Genome Consortium (ICGC)^25^. In addition, we included 43 samples from a previous proteomic analysis^26^ and 8 from the Rady Children’s Hospital Molecular Tumor Board (MTB). In total, we analyzed a retrospective cohort of WGS data of 481 tumor biopsies from 468 different patients, as well as 31 patient-derived MB models. Using DNA fingerprint analysis, we ensured that the combined cohort was not redundant, i.e., that no duplicate WGS data were present in different databases (see Methods). Clinical metadata were available for most patients and included age at diagnosis, sex, MB molecular subgroup, and survival (Supplementary Table 1). To detect ecDNA, we applied AmpliconArchitect (AA), a method and software tool that reconstructs ecDNA sequence structures from paired-end whole genome sequencing (WGS) data^1^. Among patients, ecDNA was detected in 82 (18%) cases, distributed across molecular subgroups as follows: WNT 0/22, SHH 30/112 (27%), Group 3 19/107 (18%), and Group 4 26/181 (14%) (Fig. 1a). The notable absence of ecDNA in the WNT subgroup may account for the relatively good prognosis of this subgroup^19^. Conversely, SHH tumors were significantly more likely to contain ecDNA than tumors from the other MB subgroups (χ^*2*^ =7.66, p=0.006). Among the ecDNA-amplified genes occurring in two or more samples were known or suspected MB oncogenes *MYC, MYCN, MYCL, TERT, GLI2, CCND2*^27^, and *PPM1D* (WIP1)^28^; and members of signalling pathways commonly dysregulated in cancer: DNA repair (*RAD51AP1, RAD51AP2, TOP3A* and *RAD21*); and p53 pathway inhibitors (*PPM1D, CDK6*, and *CCN4*)^29,30^ (Fig. 1b). To determine the prognostic value of ecDNA, we generated a Cox proportional hazards model parameterized on sex, age, and molecular subgroup. Patients with ecDNA (ecDNA+) were more likely to relapse (hazard ratio 2.23, p<0.005) and had shorter overall survival (hazard ratio 3.05, p<0.005) compared to patients without ecDNA (ecDNA-) (Fig. 1c,d, Supplementary Table 2).

**Figure 1:**
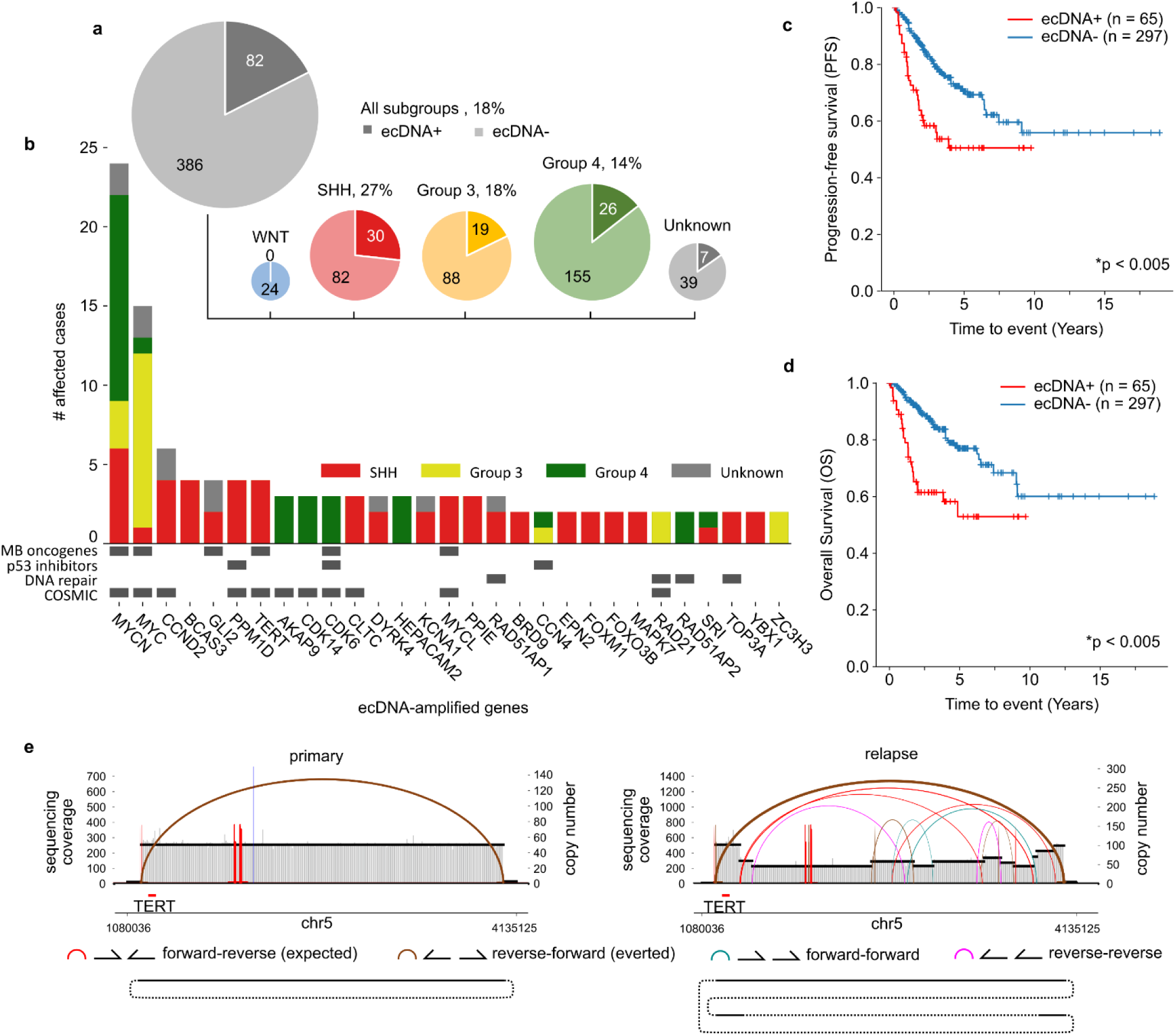
The landscape of ecDNA in medulloblastoma patient tumors. (a) Presence of ecDNA by molecular subgroup across 468 MB patient tumors. (b) Recurrently (n≥2) amplified genes on ecDNAs in this patient cohort. COSMIC: genes listed as tier 1 or 2 of the COSMIC Cancer Gene Census^31^. (c) Overall survival and (d) progression-free survival. P-values derived from Cox proportional hazards model parameterized on sex, subgroup, age, and ecDNA status. (e) Top: AmpliconArchitect visualizations of ecDNAs detected in primary and relapse biopsies of patient PT_FN4GEEFR. Bottom: visualizations showing circular sequences which can be reconstructed from WGS reads. Dotted lines indicate SV sequence rearrangement junctions.

### ecDNA sequence is rearranged during tumor progression

To investigate whether ecDNA sequences undergo structural variation during disease progression, we identified three patients with ecDNA in our cohort for whom WGS data were available from the primary and relapsed tumors. In one case (PT_CXT81GRM), there was no detectable ecDNA at diagnosis, but the relapse contained a 150x amplification of *MYC* due to ecDNA (Supplementary Fig. 1a). In the second case (PT_FN4GEEFR), the primary biopsy contained 50x ecDNA amplification of *TERT*, while the relapse tumor showed evidence of a subsequent deletion of much of the ecDNA sequence and further amplification of a smaller ecDNA subclone also containing *TERT* (Fig. 1e). In the third case (PT_XA98HG1C), the primary tumor carried an ecDNA containing regions of chr12 including *CCND2*. The primary tumor genome also contained widespread structural rearrangement along chr2q with oscillating copy number and lower-level amplification than the ecDNA, suggesting chromothripsis (Supplementary Fig. 1b,c). Chromothripsis, the catastrophic shattering of a chromosome, has recently been shown to precede ecDNA formation in cell line models^32,33^. The relapse biopsy contained 60x ecDNA amplification of *CCND2* which shared 75% of genomic regions but only 10% of breakpoints with the primary, as well as a new 60x circular amplification of an 800kb region of chr2p containing the MB oncogene *MYCN* (Supplementary Fig. 1d). Despite a small sample, our observations in these three patients suggest that ecDNA sequence rearrangement or subsequent ecDNA formation may be common events during medulloblastoma tumor progression.

### *TP53* alterations are associated with ecDNA in MB SHH tumors

The tumor suppressor protein p53 is involved in DNA damage sensing, cell cycle arrest and apoptosis, and is frequently affected by somatic mutations and pathogenic germline variants in SHH MB^21,34,35^. Moreover, SHH-subgroup MBs with inactivating *TP53* mutations are known to be associated with chromothripsis during tumor development^23^. To test whether *TP53* mutations were associated with the presence of ecDNA in SHH-subgroup MB, we accessed the somatic *TP53* mutation status available for 94 SHH MBs. Somatic *TP53* mutations were observed in 8 of 24 (33%) of SHH MB tumors with ecDNA and only 1 of 70 without ecDNA (1%), a significant enrichment (p=2.5e-5, Fisher exact test). To test whether the ecDNA+ SHH MB tumors without somatic *TP53* mutation harbor pathogenic germline *TP53* variants, we acquired *TP53* germline variant calls and annotated pathogenic variants using previously published methods^35^. As a result, we found that 12 of 23 (52%) of ecDNA+ SHH MB patients had pathogenic germline variants or somatic mutations at the *TP53* locus, compared to 2 of 69 (3%) ecDNA-SHH MB patients (p=1.3e-7, Fisher exact test).

### Patient-derived models of medulloblastoma are enriched for ecDNA relative to the clinical patient population

Patient-derived xenografts (PDX) and cell lines have been used as preclinical models for identification and testing of compounds for new targeted therapies for MB^36^. To evaluate whether ecDNA is represented in available models at a similar rate as in the clinical patient population, we analyzed WGS data derived from 27 PDX and 4 cell line models of MB for ecDNA. By subgroup, these models stratify as follows: 0 WNT, 10 SHH, 17 Group 3, and 4 Group 4 (Fig. 2a), consistent with higher engraftment rates among more aggressive molecular subgroups of other tumor types^37,38^. We find ecDNA in 19 of 31 models (61%), a significantly greater proportion than in the patient cohort (χ ^*2*^=36, p<0.005). By subgroup, ecDNA was significantly overrepresented in SHH (n=7, p=0.008, Fisher exact test) and Group 3 (n=11, p=1e-4, Fisher exact test) subgroup models relative to the patient population. *MYC* was the most frequently ecDNA-amplified gene, and all ecDNA+ Group 3 tumors contained *MYC* on the ecDNA (Fig. 2b). Overall, the relative enrichment of ecDNA in our PDX and cell line cohort suggests that the presence of ecDNA, or the overexpression of ecDNA-associated oncogenes, may predispose a tumor to successful establishment *in vitro* or *in vivo*.

**Figure 2:**
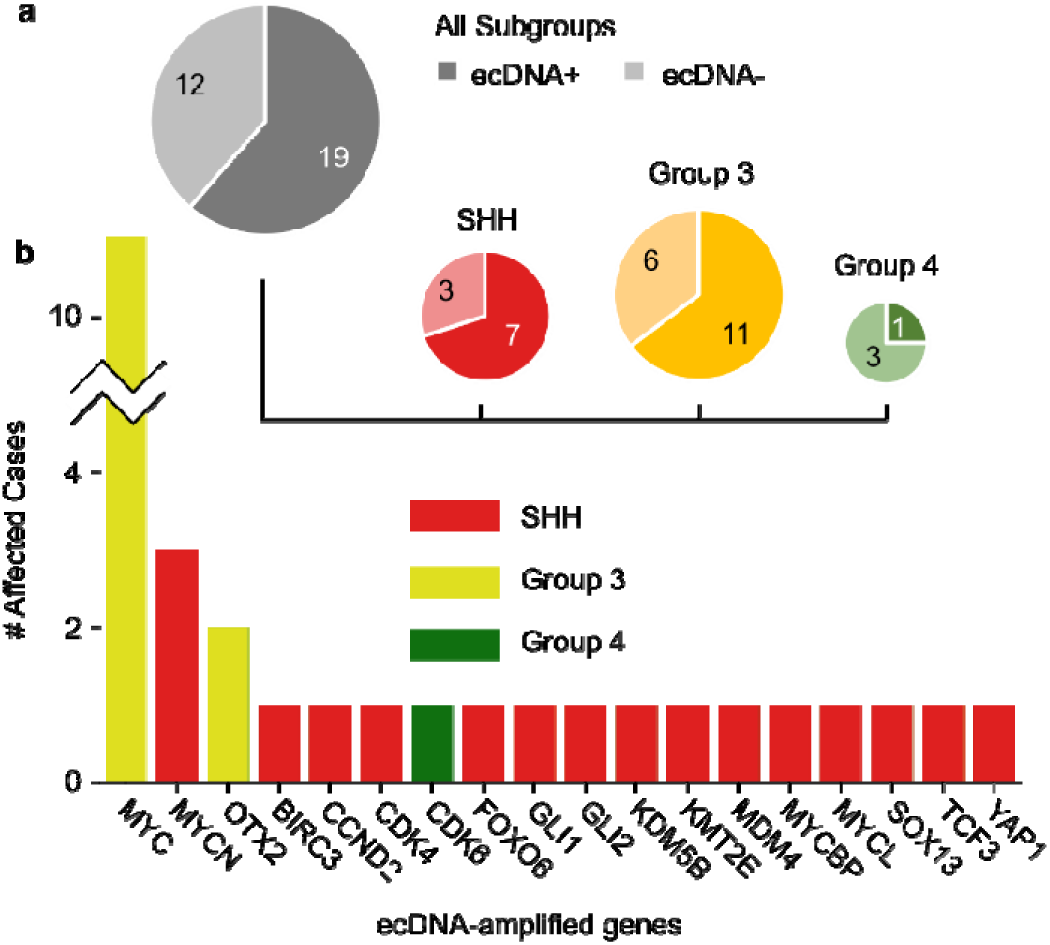
ecDNA in patient-derived model systems of MB. **(a)** Presence of ecDNA by molecular subgroup across 31 patient-derived models of MB. **(b)** Selected amplified genes on ecDNAs in MB models.

### ecDNA sequences are largely conserved in PDX models

To analyze whether ecDNA sequences are conserved during establishment and passage of PDX models, we identified 7 patients with ecDNA for whom WGS of the original human tumor (HT) and a PDX model were available (Supplementary Table 2c). To estimate the likelihood that two ecDNAs in two samples are clonally related, we measured Jaccard similarity of breakpoints and genomic regions amplified on ecDNAs (see Methods). In most (n=5, 71%) of these cases, human tumor and PDX shared at least 80% of genomic regions and two thirds of breakpoints, indicating that the sequences were largely conserved (Supplementary Table 3c). In one case (Med1911FH), we observed ecDNA originating from the same genomic locus but with heavy structural differences between the HT and the PDX (Supplementary Fig. 2a). In another case (Med211FH), ecDNA was detected only in the PDX (Supplementary Fig. 2b). We did not observe any ecDNA sequences in HTs which were subsequently lost during PDX establishment. These results underscore that PDX models can serve as faithful models of ecDNA in a human tumor, but also highlight the importance of monitoring genomic features of these model systems as they can diverge from their origin tumors^39^.

### Optical mapping establishes reference ecDNA sequences for MB models

Assembly from short paired-end sequencing cannot reliably disambiguate possible ecDNA sequences in cases where the ecDNA contains a repetitive or unmappable region, contains multiple copies of the same amplified genomic segment within a single ecDNA, or the sample contains heterogeneous subclones of the ecDNA. To establish high-confidence candidate structures for ecDNAs in MB models, we performed optical mapping (OM) of ultra-high molecular weight genomic DNA (average molecule length 342 kbp) in two MB Group 3 cell lines D425 and D458, and in the four PDX models MB002 (Group 3), Med411FH (Group 3), Icb984MB (SHH), and RCMB56 (SHH). Candidate structures were reconstructed by a combined analysis of short read WGS and *de novo* assemblies of individual OM molecules (see Methods). The average N50 (i.e., the length of the shortest contig for which 50% of the assembled length is contained in contigs of that length or longer) of the OM assemblies across all samples was 47.2Mbp (Supplementary Table 4). Reconstructed ecDNA sequence lengths ranged from 897kbp to 4.4Mbp. In all reconstructed ecDNA sequences except the cell lines D425 and D458, OM and WGS reconstruction strongly supported a single consensus ecDNA sequence (Supplementary Fig. 3); however, in each sample we assembled OM contigs which were inconsistent with the candidate reconstructed sequence, indicating sequence heterogeneity among ecDNAs present in the samples. In the Icb984MB PDX, OM assembly established a circular sequence composed of inverted tandem repeats, or head-to-head (h2h) tandem duplications, such that the reconstructed ecDNA contains 2 copies of *GLI2* (Supplementary Fig. 3d). Given that this pattern of nested h2h tandem duplication is a signature of both the breakage-fusion-bridge (BFB) cycle and tandem short template (TST) jump forms of chromosomal instability^40,41^, and that both mechanisms have been shown to precede circular amplification^32,33,42^, it is possible that the Icb984MB ecDNA arose from one or both of these mechanisms. D425 and D458 are cell lines established from the primary and relapsed tumors respectively from the same patient^43^. We therefore asked whether the ecDNAs shared sequence breakpoint junctions indicative of structural variants (SVs) present in the ancestral tumor. Although the D425 and D458 ecDNAs both contain amplifications of MB oncogenes *MYC* and *OTX2*, they share only 1 SV breakpoint junction, indicative of substantial sequence evolution after these two lines diverged from their common ancestor (Supplementary Fig. 4).

### ecDNA places oncogenes in new gene regulatory contexts

It has been shown that some MB tumors are driven by “enhancer hijacking” events, whereby somatic structural variants cause an enhancer to be rewired to amplify transcription of *GFI1* family or *PRDM6* oncogenes^20,44^. Given the extensive genomic rearrangements associated with some MB ecDNAs, we investigated whether new DNA interactions between co-amplified non-coding regulatory enhancers and oncogenes emerge on circular ecDNA, and whether such ‘enhancer rewiring’ is potentially associated with enhanced transcriptional oncogene activation. To test this hypothesis, we profiled the accessible chromatin of 25 MB tumors (11 ecDNA+, 14 ecDNA-) using ATAC-seq^45^, as well as chromatin interactions of 17 MB tumors (8 ecDNA+, 9 ecDNA-) using chromatin conformation capture (Hi-C)^46^. Consistent with previous reports^10,47^, ATAC-seq read density was markedly greater within circularized loci, even for ecDNAs with only low-level amplification as estimated by bulk WGS. Hi-C sequencing reads exhibited a similar pattern of coverage enrichment at these circularized loci (Fig. 3a).

**Figure 3:**
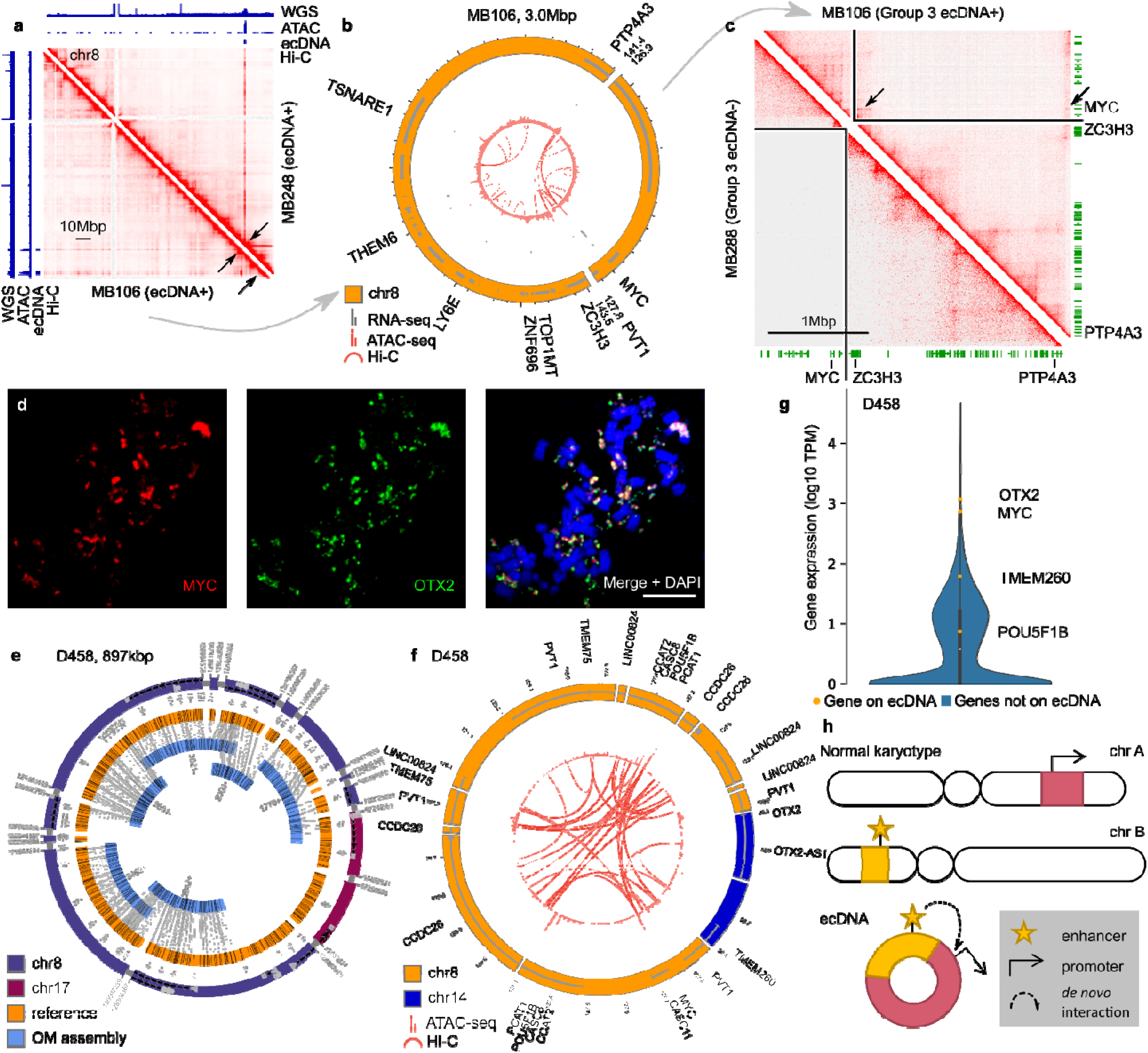
Enhancer rewiring events targeting MYC in Group 3 MB ecDNAs. **(a)** ATAC-seq and Hi-C read coverage of low-copy ecDNAs in MYC- amplified primary tumors MB248 (top right) and MB106 (bottom left). **(b)** Reconstruction of the MB106 ecDNA from WGS reads. Tracks (outer to inner): sequence (WGS), RNA-seq, ATAC-seq, Hi-C. **(c)** The Hi-C interactome of MB106 ecDNA (top right) contains enhancer- promoter interactions not visible in an unrearranged Group 3 MB tumor (bottom left). **(d)** Confocal FISH of *MYC* and *OTX2* on a metaphase spread of a D458 cell. **(e)** OM assembly of the D458 ecDNA indicates a complex ecDNA containing segments from chr8 (including MYC) and chr14 (including OTX2) with internal duplications and heterogeneous breakpoint re-use. The structure is shown in hg19 coordinates. **(f)** ATAC- seq and Hi-C interactions mapped onto the D458 amplicon reconstructed in (e). Tracks (outer to inner): sequence, ATAC-seq, Hi-C. **(g)** Gene expression of all protein-coding genes in D458. Violin plot indicates kernel density estimation of expression of n=19,178 genes not on the ecDNA. **(h)** Diagram illustration of an enhancer rewiring event as a consequence of ecDNA formation from two different chromosomes.

In half of the ecDNA+ Hi-C interaction maps (D458, MB106, MB268, and RCMB56), we observed clear evidence of aberrant chromatin interactions between genes and co-amplified enhancers which spanned DNA breakpoints (Fig. 3b,c,f, Supplementary Fig. 5). For example, in the *MYC*-amplified Group 3 MB primary tumor MB106, DNA interactions occurred between the *MYC* locus and two co-amplified enhancer regions located 13Mbp away on the linear genome (Fig. 3b). Comparing the MB106 Hi-C interactome to that of a Group 3 MB primary tumor without ecDNA, we found that these chromatin interactions are specific to the MB106 ecDNA (Fig. 3c). In the SHH MB primary tumor MB268, we identified an ecDNA amplification which includes the p53 regulator *MDM4*^48^ Supplementary Fig. 5). *MDM4* is recurrently amplified on glioblastoma ecDNAs^1^ and is a putative driver event in this tumor. In the same ecDNA, we also observed aberrant DNA interactions involving, among others, the promoter regions of the genes *LHX9, KCNT2, SNRPE*, and *KDM5B* (Supplementary Fig. 5), each of which has been previously implicated in CNS tumor development^49-52^. However, the functional significance of these co-amplified genes, enhancers, and DNA interactions in this tumor remains unclear.

In two cases, the SHH MB primary tumor biopsy RCMB56 and the Group 3 MB cell line D458, we identified interactions between DNA fragments on ecDNA from constituent loci originating from different chromosomes. D458 harbored an ecDNA amplification containing MB oncogenes *MYC* and *OTX2*, from chromosomes 8 and 14 respectively, as indicated by confocal FISH showing extrachromosomal co-localization of *OTX2* and *MYC* in D458 cells (Fig. 3d) and a candidate sequence of the D458 ecDNA assembled from WGS and OM data (Fig. 3e). Hi-C data showed potential enhancer rewiring events associating the *MYC* promoter on chromosome 8 with enhancers located on chromosome 14 (Fig. 3f). Public gene expression data from DepMap^53^ showed that *OTX2* and *MYC* were highly expressed in the D458 line (Fig. 3g). These results in multiple independent ecDNA+ tumors suggest that enhancer rewiring within ecDNA is a common event in medulloblastoma, and that tumor ecDNA amplify potential proto-oncogenes together with co-amplified enhancers, which can originate from different chromosomes (Fig. 3h).

### Distinct lineages of ecDNA co-exist in MB tumors

Although some oncogenes are recurrently amplified on ecDNA, such as *MYC* in G3 and *MYCN* in G3, G4, and SHH medulloblastoma tumors, our analyses show a large diversity of ecDNA variants across different tumors, which contain many different potential onco- or tumor-dependency genes (Fig. 1a,b). In addition, our analyses revealed a substantial diversity of ecDNA sequences within individual tumors. In total, we identified 16 patient tumors and 6 patient-derived models in which two or more ecDNAs were detected in WGS (Supplementary Table 5), each containing different potential tumor dependency genes. Assembly of available OM data supported this conclusion for the SHH PDX line Icb984MB, which harbors amplifications of MYCN and GLI2 on distinct ecDNAs (Supplementary Fig. 3c,d). In addition, WGS and OM of a PDX tumor derived from a SHH MB patient with heterozygous somatic *TP53* mutation (RCMB56) identified 2 ecDNAs, a 3.2Mbp amplicon comprising 3 regions of chr1 (RCMB56 amp1) and another 4.5Mbp amplicon comprising 23 segments originating from chr7 and chr17 (RCMB56 amp2) (Fig. 4a,b). Assembly using OM confirmed a circular structure for RCMB56 amp1 but contained a gap in RCMB56 amp2 where the amplified breakpoint junctions mapped ambiguously to peritelomeric regions (Fig. 4b). Evaluation of the WGS data derived from the tumor biopsy of patient RCMB56 confirms that these ecDNA sequences were present in the patient’s tumor and were unchanged in the PDX (Supplementary Table 3c). ecDNA copy number was estimated from bulk WGS at 20x and 10x in HT, and 30x and 25x in PDX for amp1 and amp2 respectively. Bulk ATAC-seq and Hi-C of the RCMB56 PDX revealed expected coverage enrichment mapping to both ecDNAs, as well as chromatin interactions across structural breakpoints targeting active promoters (Fig. 4c-f). These data also revealed low-level amplification of other segments of chr7 and chr17, of total length 35.2Mbp, which we hypothesized may be a third low-copy ecDNA (RCBM56 amp3) (Fig. 4f). To confirm the co-occurrence of three distinct ecDNA lineages within this tumor, we performed FISH imaging for highly transcribed genes on each amplicon: *DNTTIP2* (amp1), *KMT2E* (amp2, *KMT2E* also known as *MLL5*), and *ETV1* (amp3). FISH imaging confirmed extrachromosomal amplification of all 3 genes (Fig. 4g-i). To test whether the amplified genes are located on distinct ecDNA amplicons, we performed multi-channel FISH resulting in distinct fluorescence spots for each gene, indicating that copies of each amplified gene existed on distinct chromatin bodies (Fig. 4j-l).

**Figure 4:**
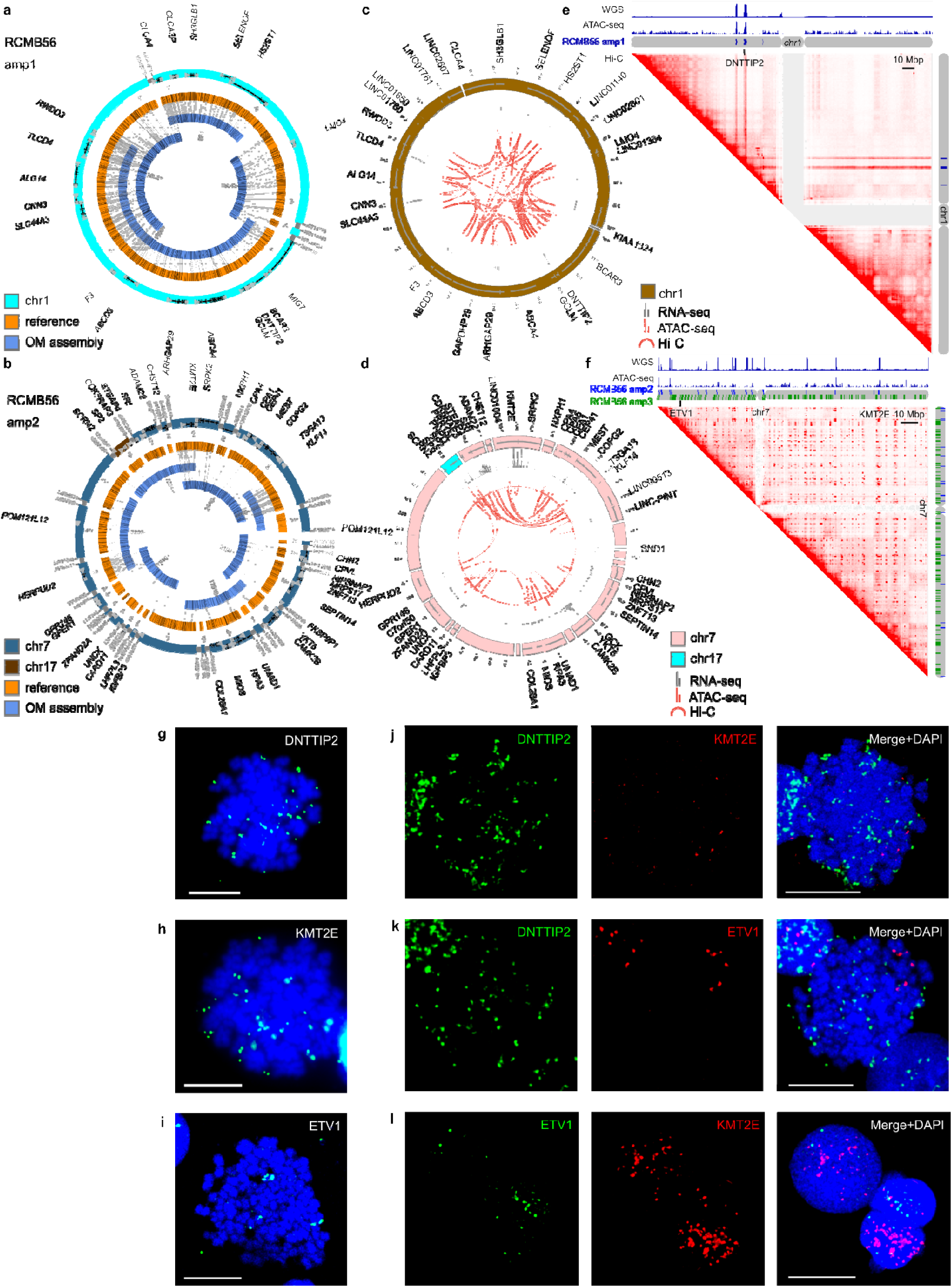
RCMB56 tissue contains multiple ecDNA lineages. **(a)** WGS and OM assembly of an ecDNA lineage (ecDNA 1) observed in the SHH MB tumor RCMB56 composed of three pieces of chr1 including DNTTIP2. The structure is shown in hg19 coordinates. The ecDNA is 3,207,166 bp long, uses all of the amplified content as detected by AA and all of the amplified breakpoints. **(b)** WGS and OM reconstruct another ecDNA lineage (ecDNA 2) 4,397,772 bp in length, with one gap mapping ambiguously to peritelomeric regions. The oscillating copy number pattern of this structure is consistent with a chromothriptic origin. **(c)** RNA-seq, ATAC-seq and Hi-C interactions mapped onto the sequence assembly of RCMB56 amp1. Chromatin interactions spanning breakpoint junctions target accessible regions at the *DNTTIP2, SH3GLB1*, and *SELENOF* gene loci. **(d)** RNA-seq, ATAC-seq and Hi-C interactions mapped onto the sequence assembly of RCMB56 amp2. Aberrant chromatin interactions spanning breakpoint junctions and targeting transcribed promoters include the *MEST, KMT2E*, and *HERPUD2* gene loci. **(e)** ATAC-seq and Hi-C read coverage is enriched at circularized regions of chr1. **(f)** ATAC-seq and Hi-C read coverage is also enriched at circularized regions of chr7 and chr17, including regions not amplified on RCMB56 amp2 and included in the proposed amp3. **(g-i)** FISH targeting amplified DNTTIP2 (amp1), KMT2E (amp2), and ETV1 (amp3) confirms extrachromosomal amplification of all 3 genes. **(j-l)** Pairwise multi- channel FISH of DNTTIP2, KMT2E and ETV1 shows non-overlapping extrachromosomal amplifications.

## DISCUSSION

Our results reveal substantial inter- and intra-tumoral heterogeneity of ecDNA lineages in particularly aggressive medulloblastoma tumors. We identify novel properties of ecDNA in medulloblastoma which may account for the unusual aggressiveness of some of these tumors. As in other cancers^3,4^, ecDNA frequently amplifies known oncogenic driver genes for MB. Our survival analysis estimates that relative to ecDNA-patients, ecDNA+ MB patients are more than twice as likely to relapse and three times as likely to die. Given this strong association with clinical outcome, it is unsurprising that ecDNA is also associated with previously known molecular markers of poor prognosis, namely *MYC* family amplification^22^ and p53 mutation^23^. Using chromosome conformation analysis (Hi-C), we find that enhancer rewiring as a result of ecDNA sequence rearrangement is a frequent event in ecDNA+ medulloblastomas. It is therefore likely that the rearrangements we observe in ecDNA sequences in MB are a consequence of selection on regulatory regions as well as genes. Further functional validation experiments are necessary to test whether the observed enhancer rewiring events contribute to transcriptional activation of co-amplified oncogenes and support tumorigenicity.

A long-standing problem in medulloblastoma oncology has been the paucity of effective targeted molecular treatments for MB, especially in relapsed cases. Recent results in glioblastoma^15^ and colon cancer^14,32,54^ cell lines implicate ecDNA in chemoresistance to targeted therapy, raising the possibility that ecDNA performs a similar role enabling MB tumors to evade treatment. The *SMO* inhibitor vismodegib, one of few targeted drugs approved for SHH medulloblastoma, is ineffective against *TP53*-mutant, *MYCN*-amplified or *GLI2*-amplified tumors^55^, each a recurrent feature of ecDNA+ MB. We find significant enrichment of ecDNA among *TP53*-mutant SHH tumors, and recurrent amplification of inhibitors of p53 among ecDNA+ tumors, adding further evidence to a possible mechanistic relationship between p53 activity and ecDNA genesis. Clarifying the mechanistic relationships between DNA repair pathway mutation, ecDNA formation and maintenance, and chemotherapy resistance may uncover new combinatorial therapies for a subset of MB patients with exceptionally poor prognoses.

## METHODS

### Medulloblastoma WGS

Paired-end whole genome sequencing (WGS) data were acquired for each of the sources described below. In total the WGS cohort comprised 468 patients, 4 cell lines and 26 PDX models (Supplementary Tables 1, 2). Unless otherwise specified, WGS was acquired for 1 tumor biosample per patient.

#### CBTN – Childrens Brain Tumor Network (114 biosamples from 101 patients)

WGS of medulloblastoma tumor biopsies were identified using the Gabriella Miller KidsFirst Data Resource center portal (https://portal.kidsfirstdrc.org/) on 29 May 2020. Patients were originally sequenced as part of the Pediatric Brain Tumor Atlas (PBTA) (https://cbtn.org/pediatric-brain-tumor-atlas). WGS data were preprocessed and aligned using the KidsFirst harmonized WGS pipeline (hg38) and subsequently analyzed using Cavatica (https://cavatica.sbgenomics.com/), on the cloud genomic analysis platform for KidsFirst genomic data. Docker containers containing fingerprint and AmpliconArchitect software were installed on the Cavatica cloud genomics platform (see Methods: “EcDNA detection and classification”).

#### St Jude (79 patients)

WGS of MB biopsies were identified using the St Jude Cloud Data Portal (https://platform.stjude.cloud/data/) on 11 February 2020. WGS was preprocessed and aligned according to internal pipelines at St Jude (hg38). Docker containers of fingerprint and AmpliconArchitect software were installed on the DNANexus cloud genomics platform (see Methods: “EcDNA detection and classification”).

#### ICGC – International Cancer Genome Consortium (237 patients)

WGS of MB tumor biopsies was identified using the ICGC Data Portal (https://dcc.icgc.org/) on 12 May 2020 and downloaded from the Collaboratory cloud genomics platform using the ICGC Score download client. These tumor genomes were previously sequenced, aligned and deposited with ICGC in other publications (GRCh37)^20,25^.

#### Archer et al. (43 patients)

All WGS data for these 43 patients were previously published elsewhere^20^. Sequencing data were aligned to human genome reference GRCh37 processed according to the best practice pipelines at the Cancer Genome Analysis group at the Broad Institute^26^.

#### Rady Childrens Hospital (8 patients)

Tumor biopsies were collected and consented for research as part of the Rady Molecular Tumor Board (MTB). Paired end reads were acquired from Rady’s Children hospital. Sequencing depth for all samples were at least 30x. Raw fastq reads were aligned to UCSC hg38 coordinates using BWA v0.7.17-r1188^56^. Reads were sorted by samtools v0.1.19^57^, marked for duplicates with Picard Tools v2.12.3, and recalibrated with GATK v3.8-1-0^58-60^.

#### Cell lines and PDX models (4 cell lines, 26 PDXs)

Low-coverage WGS for 19 PDX models and 9 corresponding origin human tumors were obtained from a previous publication^36^. An additional 6 PDX biosamples (RCMB25, RCMB32, RCMB56, RCMB57, RCMB58 and RCMB69) were contributed by Wechsler-Reya lab using similar methods to establish PDX lines^36^. Cell lines (D283, D341, D425 and D458) were contributed by Bagchi (SBP), Taylor (U. Toronto) and Wechsler-Reya (SBP) labs. Low-coverage WGS, preprocessing and alignment was performed at the UCSD IGM Genomics and Sequencing Core (hg38) for all additional samples except D283, for which no WGS was obtained.

### EcDNA detection and classification

To detect ecDNA, all samples in the WGS cohort were analyzed using AmpliconArchitect^1^ v1.2 and AmpliconClassifier^4^ v0.4.4. Briefly, the AmpliconArchitect algorithm was performed as follows. Copy number segmentation and estimation were performed using CNVkit v0.9.6^61^. Segments with copy number ≥ 4 were extracted using PrepareAA (April 2020 update) as “seed” regions. For each seed, AmpliconArchitect searches the region and nearby loci for discordant read pairs indicative of genomic structural rearrangement. Genomic segments are defined based on boundaries formed by genomic breakpoint locations (identified by discordant reads) and by modulations in genomic copy number. A breakpoint graph of the amplicon region is constructed using the CN-aware segments and the genomic breakpoints, and cyclic paths are extracted from the graph. Amplicons are classified as ecDNA, breakage-fusion-bridge, complex, linear, or no focal amplification by the heuristic-based companion script, AmpliconClassifier. Biosamples with one or more classifications of “ecDNA” were considered potentially ecDNA+, and all others were considered ecDNA-**(Suppl. Table 1c)**. We manually curated all potential ecDNA+ assembly graphs and reclassified those with inconclusive ecDNA status, which we defined as any of the following:

- Low-copy amplification (<5) AND no copy number change at discordant read breakpoints
- Cycles consisting of the repetitive region at chr5:820000 (GRCh37).

Code is available at:

- PrepareAA: https://github.com/jluebeck/PrepareAA
- AmpliconArchitect: https://github.com/virajbdeshpande/AmpliconArchitect
- AmpliconClassifier: https://github.com/jluebeck/AmpliconClassifier

The EcDNA-status of the D283 cell line was not determined computationally by WGS, but by copy number analysis of DNA methylation, FISH (see Methods: “FISH”), and analysis of OM data.

### Fingerprinting analysis

To uniquely identify WGS from each patient, we counted reference and alternate allele frequencies at 1000 variable non-pathogenic SNP locations in the human genome according to the 1000 Genomes project^62^, and performed pairwise Pearson correlation between all WGS samples. Biospecimens originating from the same patient tumor (eg., primary/relapse or origin/PDX pairs) were readily distinguishable by high correlation across these sites (r > 0.80). We identified one case in our cohort in which 2 tumor biosamples had highly correlated fingerprints: MDT-AP-1217.bam and ICGC_MB127.bam. We arbitrarily removed ICGC_MB127 from the patient cohort.

### Patient metadata, survival, and medulloblastoma subgroup annotation

Where available, patient samples and models were assigned metadata annotations including age, sex, survival, and MB subgroup based on previously published annotations of the same tumor or model^20,26,35,36,52,63,64^. Sample metadata are also available in some cases from the respective cloud genomics data platform: https://dcc.icgc.org/ (ICGC), https://pedcbioportal.kidsfirstdrc.org/ and https://portal.kidsfirstdrc.org/ (CBTN), and https://pecan.stjude.cloud/ (St Jude). Where primary sources disagreed on a metadata value, that value was reassigned to NA. Patient tumors from the CBTN were assigned molecular subgroups based on consensus of 2 molecular classifiers using RSEM-normalized FPKM data: MM2S^65^ and the D3b medulloblastoma classifier at the Children’s Hospital of Philadelphia (https://github.com/d3b-center/medullo-classifier-package). To determine molecular subgroup of PDX samples, we generated or obtained from a previous publication^36^ DNA methylation profiles (Illumina 450k or EPIC) and classified samples by molecular subgroup according to the DKFZ brain tumor methylation classifier (https://www.molecularneuropathology.org/mnp)^20^.

### *TP53* mutation annotation

#### Somatic mutations

Somatic *TP53* mutation information for the ICGC and CBTN cohorts was acquired from a previous publication^35^ and from the ICGC and CBTN data portals (see Methods: “Patient metadata, survival, and subgroup annotation”. Somatic *TP53* mutation information for the St. Jude cohort was extracted from the standard internal St. Jude variant calling pipeline^24^. Somatic mutations were only considered which were protein-coding and missense, nonsense, insertion or deletion.

#### Germline variants

Whole-genome sequencing (WGS) GVCF files were downloaded from the ICGC data portal (https://dcc.icgc.org/), the KidsFirst data portal (https://portal.kidsfirstdrc.org/dashboard) and DNAnexus for St. Jude Pediatric Cancer Genome Project (PCPG). GVCF files were merged with GLnexus^66^ and converted to PLINK format for analysis of ICGC, PCPG and KidsFirst genotypes. PCPG genotypes were converted to hg19 coordinates using liftover. Variants from *TP53* genomic locus (hg19:chr17:7571739-759080) were extracted and annotated with REVEL^67^, CADD^68^, ClinVar (accessed June 2021) and Variant Effect Predictor (VEP)^69^. REVEL GRCh38 scores were downloaded from https://sites.google.com/site/revelgenomics/ (date accessed: 03/05/2021). CADDv1.6 scores were downloaded from https://cadd.gs.washington.edu/info (date accessed: 03/23/2020). VEP scores were calculated with http://grch37.ensembl.org/index.html release 104 (date accessed: 04/27/21). Clinvar scores were obtained from https://ftp.ncbi.nlm.nih.gov/pub/clinvar/vcf_GRCh37/ (Date accessed: 04/27/21). VEP variants considered pathogenic included “frameshift” and “splice” variants. ClinVar annotations considered pathogenic included “frameshift”, “stop”, “splice”, and “deletion”, and whose clinical significance were “pathogenic” or “likely pathogenic”. CADD “pathogenic” variants had a CADD score of at least 10 (Phred score). REVEL “pathogenic” variants had a REVEL score of at least 0.5. Only variants with minor allele frequency (MAF) less than 5%, according to the gnomAD r2.1.1 database, were analyzed^70^.

### Optical mapping data collection and processing

Ultra-high molecular weight (UHMW) DNA was extracted from frozen cells preserved in DMSO following the manufacturer’s protocols (Bionano Genomics, USA). Cells were digested with Proteinase K and RNAse A. DNA was precipitated with isopropanol and bound with nanobind magnetic disks. Bound UHMW DNA was resuspended in the elution buffer and quantified with Qubit dsDNA assay kits (ThermoFisher Scientific).

DNA labeling was performed following manufacturer’s protocols (Bionano Genomics, USA). Standard Direct Labeling Enzyme 1 (DLE-1) reactions were carried out using 750 ng of purified UHMW DNA. The fluorescently labeled DNA molecules were imaged sequentially across nanochannels on a Saphyr instrument. A genome coverage of at least 400x was achieved for all samples.

*De novo* assemblies of the samples were performed with Bionano’s *De Novo* assembly Pipeline (DNP) using standard haplotype aware arguments (Bionano Solve v3.6). With the Overlap-Layout-Consensus paradigm, pairwise comparison of DNA molecules was used to create a layout overlap graph, which was then used to generate the initial consensus genome maps. By realigning molecules to the genome maps (P value cutoff of <10^−12^) and by using only the best matched molecules, a refinement step was done to refine the label positions on the genome maps and to remove chimeric joins. Next, during an extension step, the software aligned molecules to genome maps (P<10^−12^), and extended the maps based on the molecules aligning past the map ends. Overlapping genome maps were then merged (P<10^−16^). These extension and merge steps were repeated five times before a final refinement (P<10^−12^) was applied to “finish” all genome maps.

### ecDNA reconstruction with OM data

We used an ecDNA reconstruction strategy which incorporated the short-read derived CN-aware breakpoint graph generated by AA^1^ with OM contigs generated by the Bionano *de novo* assembly Pipeline, and in RCMB56 we utilized contigs from both the Bionano DNP as well as the Rare Variant Pipeline (RVP).

We used AmpliconReconstructor^71^ (AR) v1.01 to scaffold together individual breakpoint graph segments using the collection of OM contigs. We ran AR with the --noConnect flag set and otherwise default settings. A subset of informative contigs with alignments to multiple graph segments as well as a breakpoint junction were then selected for subsequent scaffolding by AR. For exploration of unaligned regions of OM contigs used in the reconstructions, we utilized the OM alignment tool FaNDOM^72^ v0.2 (default settings). FaNDOM was used to identify the loose ends of the RCMB56 ecDNA2.

A number of samples were amenable to AR-based reconstruction methods (Icb984MB ecDNA 1 & 2, MB002, Med411FH, RCMB56 ecDNA 1), however D425, D458 and RCMB56 ecDNA 2 required more manual intervention. Due to the fractured nature of the breakpoint graph in D425 and RCMB56 ecDNA 2, we searched for CN-aware paths in the AA breakpoint graph (using the plausible_paths.py script from PrepareAA), then converted these to *in silico* OM sequences and aligned paths to OM contigs directly using AR’s SegAligner.

Due to heterogeneity suggested by the OM contigs in D458, we instead utilized a reconstruction strategy involving alignment of individual OM molecules to the putative D458 cyclic path suggested from short reads alone. After converting the path to an *in silico* OM sequence, we aligned OM molecules (instead of contigs) directly to structure to validate the breakpoint junctions.

### ATAC-seq

#### Archer et al. samples

At least 25mg of pulverized frozen tissue from samples previously included in proteomics analysis^26^ were sequenced by ATAC-seq according to an established protocol for bulk ATAC-seq of frozen neuronal cells^73^. Reads were aligned, deduplicated and preprocessed according to ENCODE best practices. Samples included in subsequent analyses passed the following quality control thresholds: PCR bottleneck coefficient 1 (PBC1) > 0.7, PCR bottleneck coefficient 2 (PBC2) > 3, non-redundant read fraction (NRF) > 0.8 and TSS enrichment > 2.6. These samples had at least 13 million uniquely mapped single-end reads (GRCh37). Accessible chromatin regions were identified using MACS2 v2.1.2^74^ using Benjamini-Hochberg corrected p-value threshold < 0.05.

#### RCMB56 and D458

Frozen dissociated cells were ATAC-sequenced at ActiveMotif, Inc. (San Diego, CA). Briefly, ∼100K cells were permeabilized and transposase loaded with sequencing adapters was added. Sequencing was performed on Illumina NextSeq 500. Reads were aligned, deduplicated and preprocessed according to ENCODE best practices. Samples had PBC1 > 0.9, PBC2 > 20, NRF > 0.5 and at least 30 million uniquely mapped paired-end reads (hg38). Accessible chromatin regions were identified using MACS2 v2.1.2^74^ using Benjamini-Hochberg corrected p-value threshold < 0.05.

### Chromosome conformation capture

#### Archer et al. samples (except MB248, MB275)

Hi-C on frozen tumor tissue sample was carried out using protocols previously described for tissue Hi-C experiments^75^. In brief, frozen tissues are pulverized using a mortar and pestle kept cold on a bed of dry ice into a fine powder. The tissue powder was then transferred to a 15mL conical tube containing 5mLs of DPBS and fixed with 2% formaldehyde for 10 minutes. The fixation was quenched by addition of 0.2M Glycine. The fixed tissue was pelleted by centrifugation, washed 1x with DPBS, and then flash frozen until ready for further processing. For Hi-C experiments, the fixed frozen tissue pellets were first resuspended in 3mLs of lysis buffer (10mM Tris-HCl pH 8.0, 5mM CaCl_2_, 3mM MgAc, 2mM EDTA, 0.2mM EGTA, 1mM DTT, 0.1mM PMSF, 1X Complete Protease Inhibitors). The sample was transferred to an M-tube and dissociated using a GentleMACS Tissue dissociator (Miltenyi) using the “Protein M-tube” setting. The sample was removed from the M-tube into a 50mL conical. The M-tube was washed with 3mLs of lysis buffer with 0.4% Triton X-100 added, and this wash was combined with the original 3mLs of sample for a total volume of 6mLs with final concentration of 0.2% Triton X-100. The sample was then passed through a 40µM cell strainer. The strainer was washed with an additional 2mLs of lysis buffer with 0.2% Triton X-100. The sample was then centrifuged and washed with 1mL of lysis buffer with 0.2% Triton X-100. After centrifugation, the sample was resuspended in 0.5% SDS and processed with previously described in situ Hi-C method^46^ using the MboI enzyme. Libraries were prepared using the Illumina TruSeq LT sequencing adaptors. Initial QC sequencing was first performed on a MiSeq to assess library quality, and if sufficient, was subject to production scale sequencing on the HiSeq X or NovaSeq platform, respectively.

#### D458, RCMB56, MB248 and MB275

Hi-C experiments were carried out by Arima Genomics, Inc (San Diego, CA) using the Arima-HiC Kit protocol with MboI restriction enzyme. Subsequently, Illumina-compatible sequencing libraries were prepared by shearing the proximally ligated DNA and then size-selecting DNA fragments using SPRI beads. The size-selected fragments containing ligation junctions were enriched using Enrichment Beads (provided in the Arima-HiC Kit), and converted into Illumina-compatible sequencing libraries using the Swift Accel-NGS 2S Plus kit (P/N: 21024) reagents. After adapter ligation, DNA was PCR amplified and purified using SPRI beads. The purified DNA underwent standard QC (qPCR and Bioanalyzer) and sequenced on the NovaSeq following manufacturer’s protocols.

### Hi-C data processing

Hi-C reads were trimmed using Trimmomatic 0.39^76^ and aligned to the hg38 human genome reference using the HiC-Pro toolkit v2.11.3-beta using default parameters^77^. Visualization and contact normalization was performed with JuiceBox v1.11.08^78^ and the Knight-Ruiz algorithm^79^. Chromatin interactions were called using Juicer Tools GPU HiCCUPS v1.22.01^80^ using fdr threshold 0.2 and default recommended parameters^46^ for Hi-C. From visual inspection, we found that HiCCUPS correctly annotated interactions mapping to ecDNA, except for locus pairs mapping within ∼50kb of a structural rearrangement. Due to these technical challenges, chromatin interactions described herein were manually curated based on HiCCUPS interaction calls.

### Animals

NOD-SCID IL2R*γ* null (NSG) mice used for intracranial human tumor transplantation were purchased from The Jackson Laboratory (#005557). Mice were bred and maintained in the animal facilities at the Sanford Consortium for Regenerative Medicine. All experiments were performed in accordance with national guidelines and regulations, and with the approval of the animal care and use committees at the Sanford Burnham Prebys Medical Discovery Institute and University of California San Diego (San Diego, CA, USA).

### Establishment and maintenance of PDX RCMB56

PDX RCMB56 was established by implanting 0.5-1×10^6^ dissociated patient tumor cells directly into the cerebellum of NSG mice. Subsequent tumors were harvested from mice, dissociated and reimplanted into new NSG mice without *in vitro* passaging. *Ex vivo* experiments were performed with PDX RCMB56 cells of *in vivo* passage 1 (p1) or greater.

### Metaphase spreads

Cell lines were enriched for metaphases by addition of KaryoMAX (Gibco) at 0.1µg/mL for between 2h. – overnight (0.02µg/mL overnight for dissociated PDX cells). Single cell suspensions were then incubated with 75mM KCl for 8-15 minutes at 37°C. Cells were then fixed by carnoy fixative (3:1 methanol:acetic acid) and washed in fixative 3 times. Cells were then dropped onto humidified slides.

### FISH

Slides containing fixed cells were briefly equilibrated in 2X SSC, followed by dehydration in 70%, 85%, and 100% EtOH for 2 minutes each. FISH probes (Empire Genomics) diluted in hybridization buffer (Empire Genomics) were applied to slides and covered with a coverslip. Slides were denatured at 72°C for 1-2 minutes and hybridized overnight at 37°C. The slide was then washed with 0.4x SSC, then 2x SSC-0.1% Tween 20. DAPI was added before washing again and mounting with Prolong Gold.

### Microscopy

Conventional fluorescence microscopy was performed using an Olympus BX43 microscope, and images were acquired with a QiClick cooled camera. Confocal microscopy was performed using a Leica SP8 microscope with lightning deconvolution and white light laser (UCSD School of Medicine Microscopy Core). Excitation wavelengths for multiple color FISH images were set manually based on the optimal wavelength for the individual probes, with care taken to minimize crosstalk between channels. ImageJ was used to uniformly edit and crop images.

### Pairwise similarity of ecDNAs sequences

We compared overlapping focal amplifications to quantify amplicon similarity by quantifying the relative degrees of shared overlap in genomic coordinates and in SV breakpoint location. These calculations are implemented into the amplicon_similarity.py script, available in the AmpliconClassifier repository (https://github.com/jluebeck/AmpliconClassifier).

We defined two measurements of similarity based on Jaccard indexes. First, JaccardGenomicSegment similarity, which is a Jaccard index computed on two sets formed by the coordinate ranges of the genomic intervals comprising two focal amplifications. Second, JaccardBreakpoint similarity, which is a Jaccard index computed on two sets formed by the locations of SV breakpoint junctions in the two focal amplifications. Two SV breakpoint junctions were determined to be the same if the total absolute difference between the measured genomic endpoints of the junction was less than 250bp.

The amplicon similarity script supports comparison of amplicons both globally for all amplicon regions, but also can be run in a restricted mode which limits the comparison to specific regions of the genome. Furthermore, as AmpliconArchitect may include flanking regions which are not focally amplified as part of the amplification itself, the amplicon similarity script filters from the calculation regions that are not focally amplified (CN < 4.5 default), and we also redundantly filter regions that are also present in the low-complexity or low-mappability database used by AmpliconArchitect.

### Statistical methods

Statistical test, test statistic and p-values are indicated where appropriate in the main text. Categorical associations were established using the chi-squared test of independence if N>5 for all categories, and the Fisher exact test otherwise. For both tests, the python package scipy.stats v1.5.3 implementation was used^81^. Multiple hypothesis corrections were performed using the Benjamini-Hochberg correction method implemented in statsmodels v0.12.0^82^. Kaplan-Meier and Cox Proportional Hazards analyses were performed with Lifelines v0.21.0^83^.

## Supporting information

Supplementary Table 1 Patient Cohort

Supplementary Table 2 Cox Proportional Hazards

Supplementary Table 3 Model Cohort

Supplementary Table 4 OM Statistics

Supplementary Table 5 Multiple ecDNA

## ACKNOWLEDGMENTS

This work is supported by a Hannah’s Heroes St. Baldrick’s Scholar Award (LC) and funding from the NIH National Institute of Neurological Disorders and Stroke Institute R21 NS116455 (LC) and R21 NS120075 (LC), the NIH National Cancer Institute R01 CA159859, U01 CA184898 (JPM) and U24 CA264379 (VB and JPM), the NIH National Institute of General Medical Sciences R01 GM074024 (JPM) and R01 GM114362 (VB and JPM), the NIH National Library of Medicine T15 LM011271 (OSC), and a Moores Cancer Center Pilot Grant (LC, VB, JPM, and PSM). Microscopy work was supported by funding from the NIH National Institute of Neurological Disorders and Stroke NS047101 (UCSD Microscopy Core). This work used the Extreme Science and Engineering Discovery Environment (XSEDE), which is supported by National Science Foundation grant number ACI-1548562. DNA methylation array analysis was conducted at the IGM Genomics Center, University of California, San Diego, La Jolla, CA (MCC grant # P30CA023100). This research was conducted using data made available by The Children’s Brain Tumor Network (formerly the Children’s Brain Tumor Tissue Consortium).

In addition, we thank Iris Anne Reyes, Jinghui Zhang, Clay McLeod, and Adam Resnick for facilitating data access; Michael Reich, Mahidhar Tatineni and Xinlian Zhang for computational support; Jim Olson, Anindya Bagchi, Jae Cho and Xiao-Nan Li for providing biosamples; and Alex Wenzel and Michael Chapman for helpful scientific discussion.

## CONTRIBUTIONS

O.S.C., J.L., J.P.M. and L.C. prepared the manuscript and figures. S.W., A.T., S.C., M.A., J.R.C., N.C., M.L., D.M.M., S.L.P., J.D., R.H.S. and E.F. performed sample preparation and experimental analysis. J.D.L. and R.W.R. contributed PDX mouse models. J.T.L., I.T.L.W., and P.S.M. performed all microscopy. M.P., S.Q.W., Y.L., S.S., J.R., S.R.D., E.J., H.C. and V.B. contributed computational data analyses. O.S.C., J.P.M., and L.C. designed the study and J.P.M. and L.C. co-supervised the project.

## ETHICS DECLARATIONS

The authors declare the following competing interests:

V.B. is a co-founder, consultant, SAB member and has equity interest in Boundless Bio, inc. and Abterra, Inc. The terms of this arrangement have been reviewed and approved by the University of California, San Diego in accordance with its conflict of interest policies.

## BIOLOGICAL MATERIAL AVAILABILITY

PDX and cell line materials used in this study were previously described^36,43^. Patient tumor material used in this study are depleted and therefore not available.

## DATA AVAILABILITY

Whole genome sequencing data analyzed in this work are under controlled access, but are available from the following sources upon reasonable request:

- ICGC and Archer patient cohorts: International Cancer Genome Consortium (https://dcc.icgc.org/)
- CBTN patient cohort: Kids First Data Resource Center (https://kidsfirstdrc.org/)
- St Jude patient cohort: St Jude Cloud (https://www.stjude.cloud/)
- MB cell line and PDX models: data available upon reasonable request.

RNA-seq data will be available from NCBI Gene Expression Omnibus (GEO). ATAC-seq and Hi-C data will be available from the NCBI Sequence Read Archive (SRA). Accession numbers will be provided in the final publication.

## CODE AVAILABILITY

Code for the AmpliconArchitect family of software tools is available from the following repositories:

Source code for all other computational analyses is available upon reasonable request.

## SUPPLEMENTARY FIGURES

**Supplementary Figure 1:**
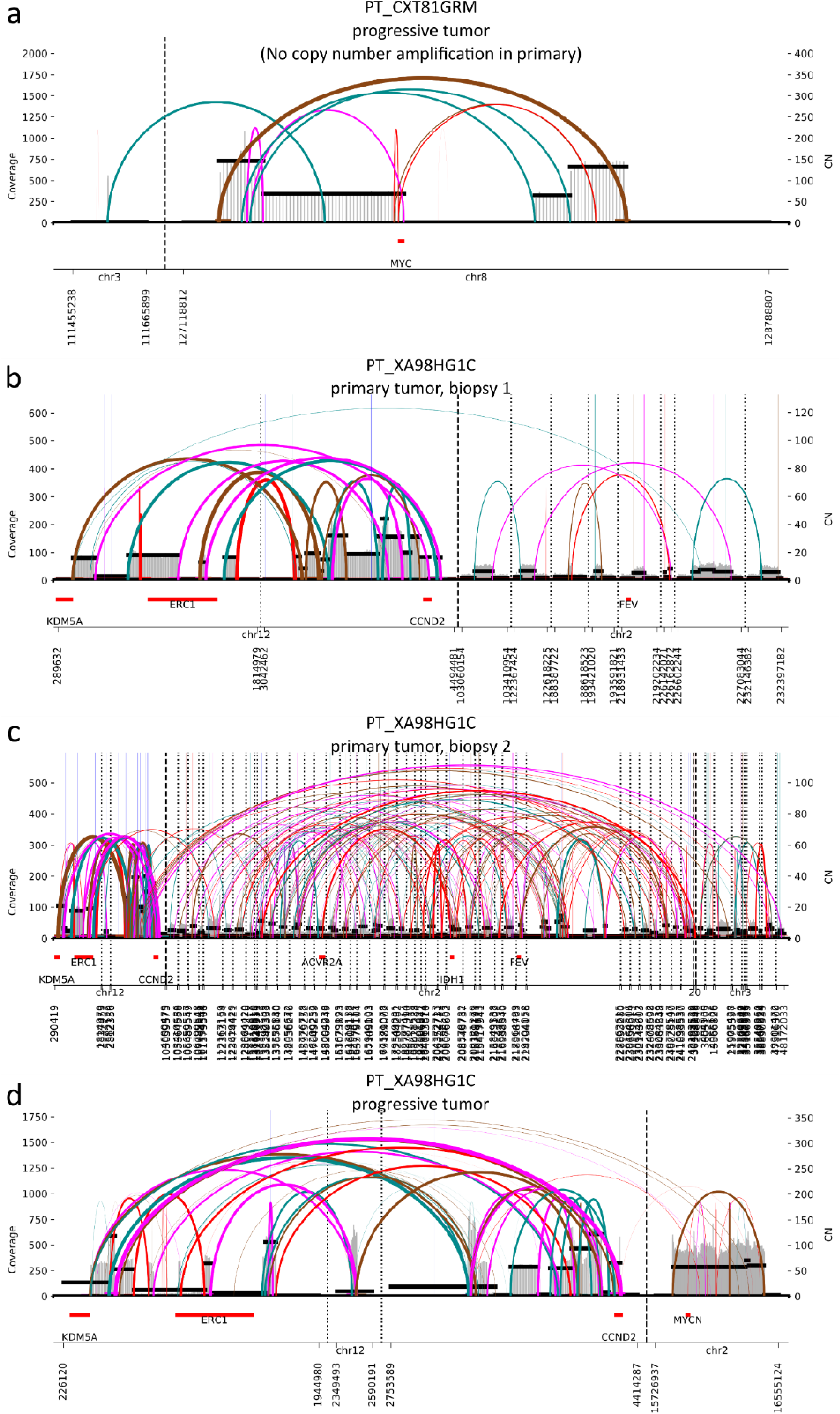
AmpliconArchitect identifies structural mutation of ecDNA between primary and relapsed patient biopsies. (**a**) AmpliconArchitect (AA) identifies a cyclic amplification of a segment derived from chr8 containing MYC in WGS from the biopsy of progressive medulloblastoma from PT_CXT81GRM. Copy number of MYC was estimated by AA at 63. Primary patient tumor was classified as SHH subgroup from RNA expression data (see Methods). Pathogenic somatic TP53 mutation was found in the progressive but not the primary tumor. Genomic coordinates are relative to the hg38 reference assembly. No amplification was detected at this locus in the primary tumor. (**b-d**) AmpliconArchitect identifies cyclic amplifications of overlapping segments of chr12 in WGS of two distinct primary biopsies (**b**, biopsy 1; **c**, biopsy 2) and one progressive tumor biopsy from patient PT_XA98HG1C, including the same fusion event CCND2-NINJ2. Primary patient tumor was classified as SHH subgroup from RNA expression data (see Methods). Copy number of the fusion gene is estimated as 16 in the primary and 109 in the progressive. The progressive sample also contained circular amplification of chr2 including the MB oncogene MYCN at estimated copy number 59.

**Supplementary Figure 2:**
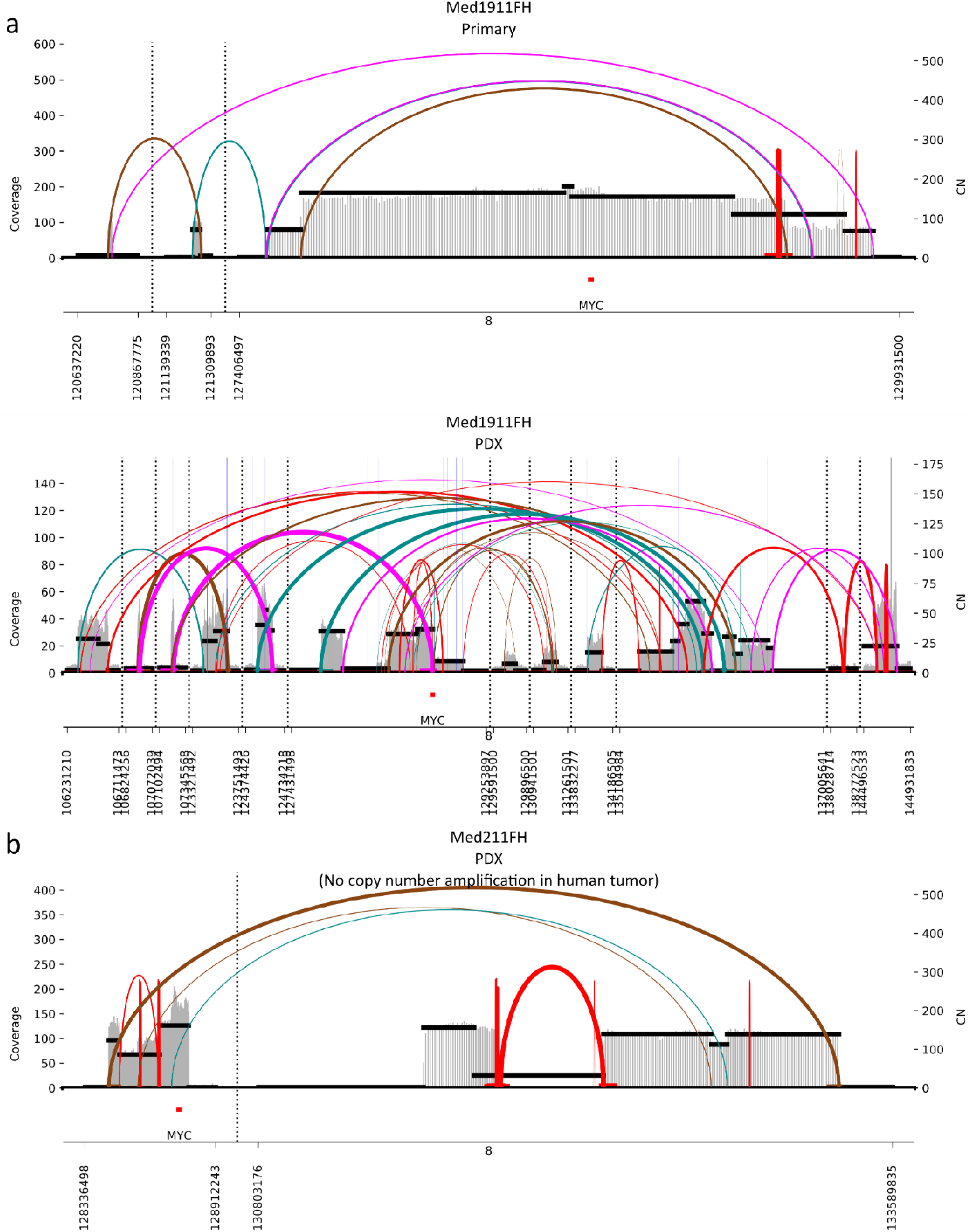
AmpliconArchitect identifies structural mutation of ecDNA between PDX models and their origin human tumors. **(a)** AmpliconArchitect identifies circular amplifications of *MYC* in the human tumor (HT) and PDX of the group 3 MB model Med1911FH. No breakpoint junctions are shared between the two amplicons. (**b**) AmpliconArchitect identifies a high-copy *MYC* amplification in the PDX, but not the origin tumor, of Med211FH. Assembly from discordant paired-end reads was unable to reconstruct a cyclic structure but the amplification is predicted to be ecDNA due to high-copy focal amplification.

**Supplementary Figure 3:**
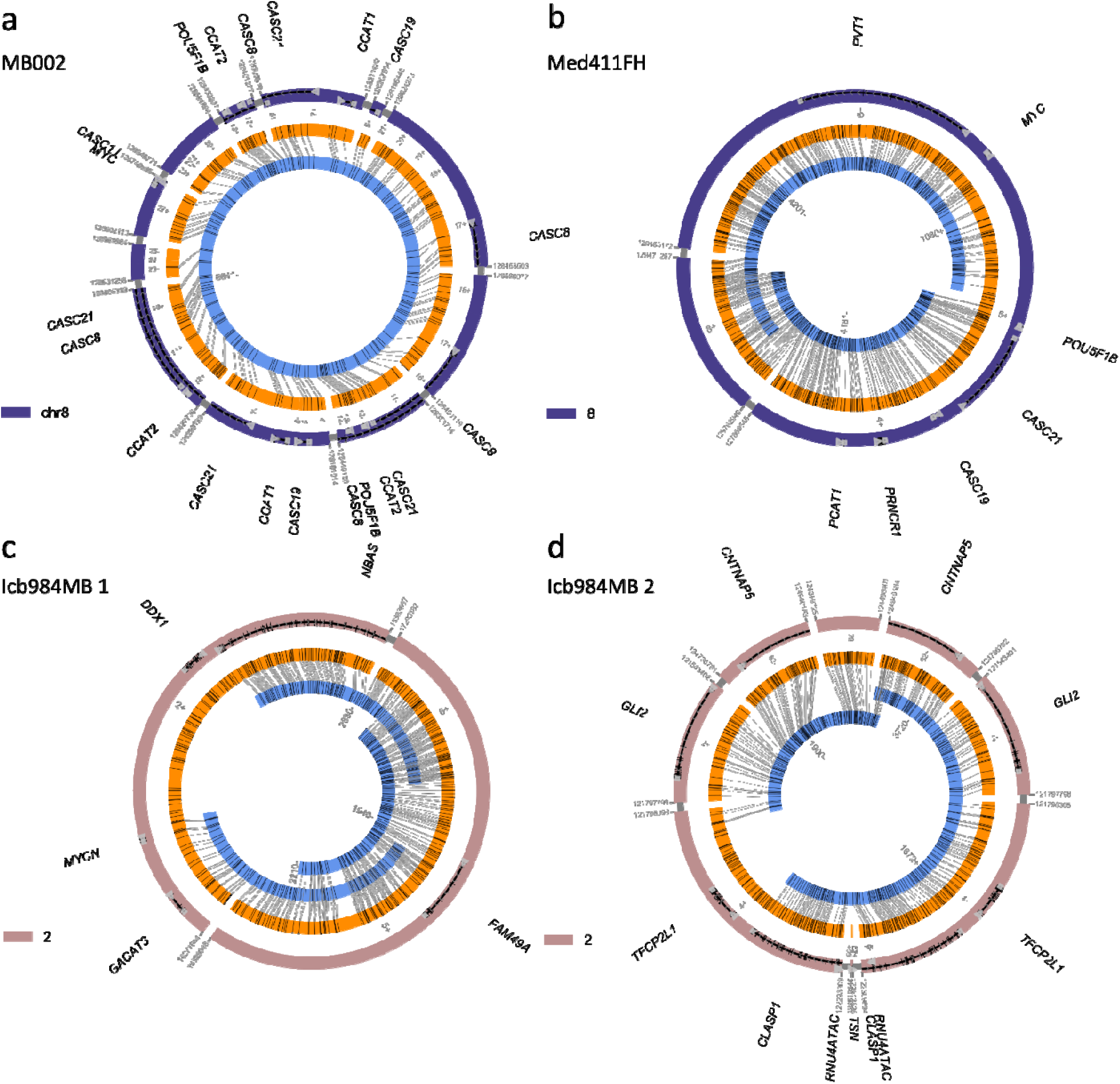
Whole-genome sequencing and optical mapping establish high-confidence reference sequences for MB PDX models. (**a**) Assembly from optical mapping (OM) of Group 3 PDX line MB002 identifies a circular contig containing *MYC*. The structure is shown in hg19 coordinates. The ecDNA is 906,496 bp long, contains 86.5% of the amplified content (CN > 10), and uses 11 of 12 breakpoints from amplified regions identified from paired-end sequencing. (**b**) A trivial ecDNA structure derived from chr8 (containing MYC and intact PVT1) with a 6kbp deletion, observed in the G3 MB PDX line Med411FH. The structure is shown in hg19 coordinates. The ecDNA is 1,897,314 bp long, uses all of the amplified content (CN >10) and both breakpoints in amplified regions identified by PE sequencing. The OM scaffolds and resulting structure were generated using AR. (**c**) A trivial ecDNA structure derived from chr2 containing MYCN. The ecDNA is 1,841,608 bp long and uses 97% of amplified content (CN > 50). The structure is shown in hg19 coordinates. The gap region in the AA amplicon overlaps a known gap region of reference hg19. The OM scaffolds and resulting structure were generated using AR. (**d**) An agglomerated complex ecDNA structure derived from chr2 containing GLI2. The agglomerated structure contains 1,099,546 bp of amplified genomic content (87.3% of the total amplified content with CN >10 as suggested by AA) scaffolded into a structure containing duplications and a ‘head-to-head’ agglomeration pattern >3 Mbp in length. The OM scaffolds and resulting structure were generated using AR. A 168kbp gap in the reconstruction appears in contig 1900, however the resulting reconstruction remains circular.

**Supplementary Figure 4:**
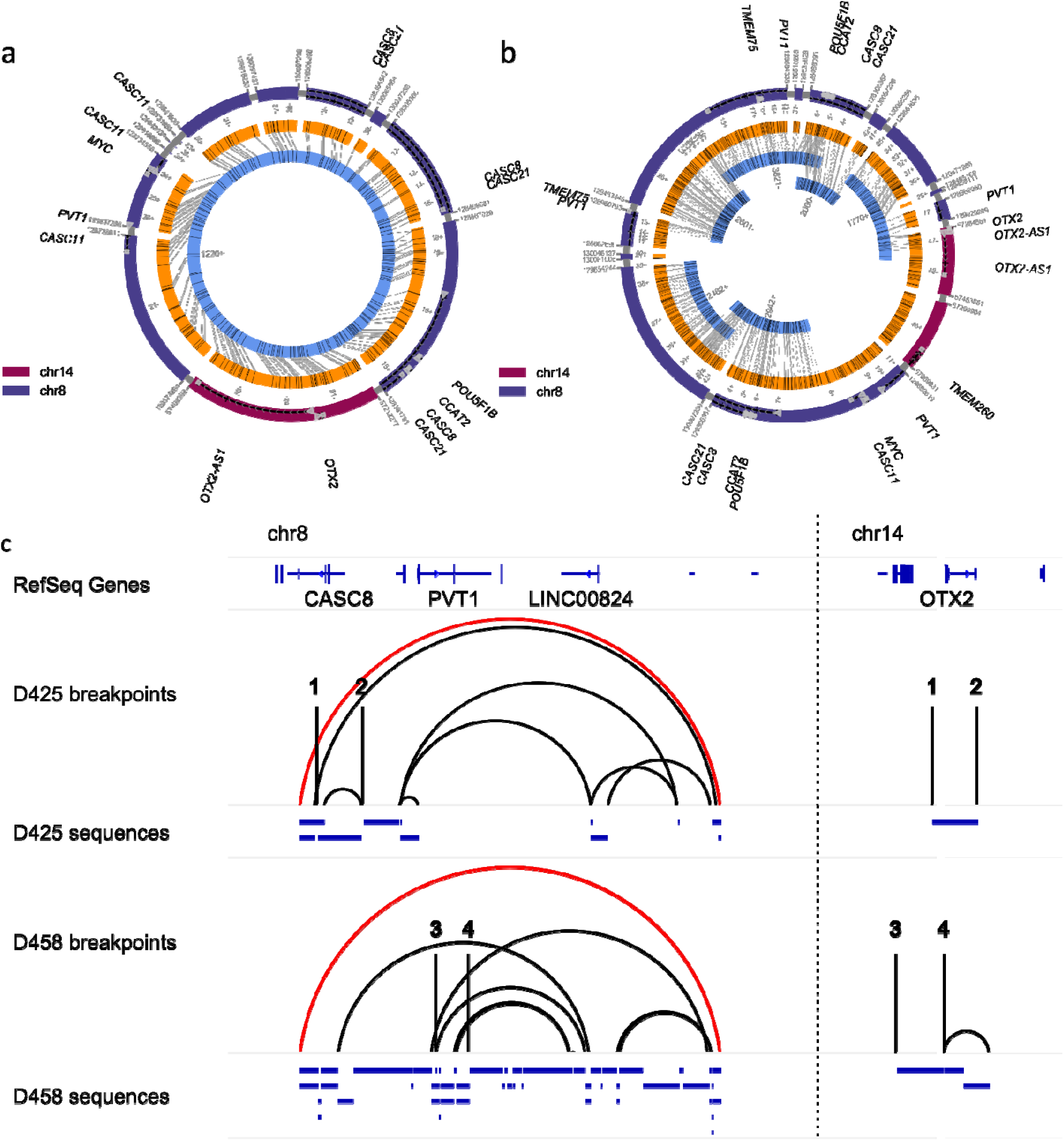
The ecDNAs of Group 3 MB cell lines D425 and D458 contain a shared ancestral breakpoint junction. (**a**) A complex ecDNA containing segments from chr8 (including MYC) and chr14 (including OTX2) with internal duplications and heterogeneous breakpoint re-use. The structure is shown in hg19 coordinates. The ecDNA is 896,524 bp long, uses all of the amplified content (CN> 50) as detected by AA and uses 9 of 12 breakpoints from amplified regions. Due to the highly segmented nature of the breakpoint graph, the reconstruction was performed by searching AA-supported paths and aligning the paths to contigs using SegAligner. We identified a circular Bionano contig which supported the circular ecDNA structure. (**b**) D458 contains a complex ecDNA comprising segments of chr8 (including MYC) and chr14 (including OTX2) with internal duplications and heterogeneous breakpoint re-use. The structure is shown in hg19 coordinates. The ecDNA is 2,575,873 bp long, uses all of the amplified genomic content (CN > 10) as detected by AA and uses 12 of 21 breakpoints from amplified regions. The OM scaffolds we identified with AR were highly heterogeneous, and the structure we found demonstrates some partially aligned OM contigs to a cyclic structure identified using AA breakpoints as no single structure could be reconciled from both sources of data. (**c**) Sequences and breakpoint junctions in the reconstructions in (a) and (b) mapped onto the hg19 reference sequence. Numbered breakpoints 1-4 indicate either end of a trans- chromosomal SV breakpoint. The two ecDNAs share much of the same sequence and a single breakpoint, highlighted in red, consistent with their shared ancestral origin in the primary and relapsed tumors of a Group 3 MB patient.

**Supplementary Figure 5:**
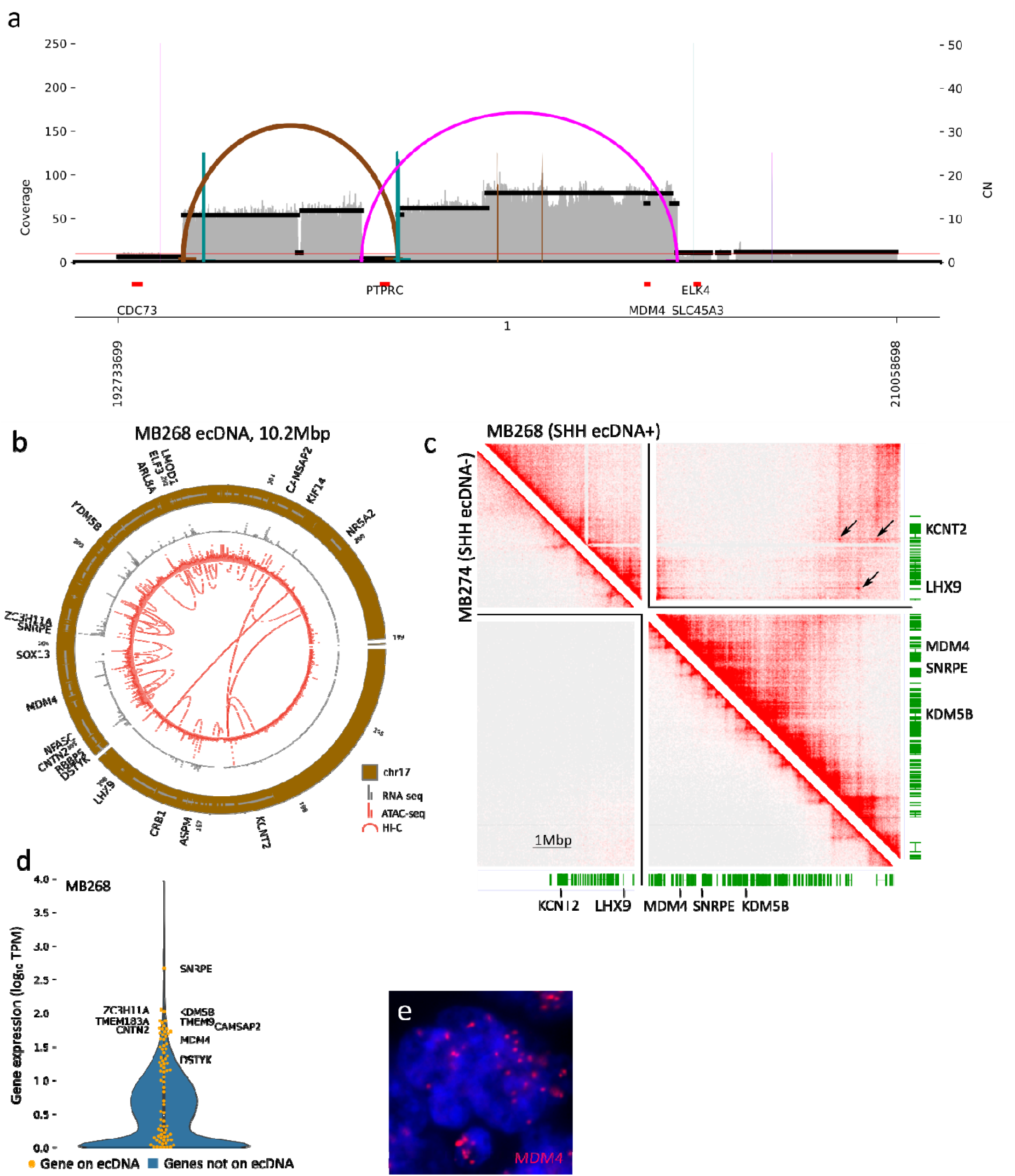
The epigenetic profile of the MB268 ecDNA. (**a**) AA resolves a circular structure composed of 3 segments of chr1 from short paired-end reads derived from the SHH MB tumor MB268. (**b**) RNA-seq, ATAC-seq and Hi-C interactions mapped onto the ecDNA sequence. Amplified oncogenes include *MDM4*, a p53 inhibitor frequently amplified on ecDNAs of cancers of various types. Chromatin interactions spanning breakpoints target accessible regions at the *LHX9* and *KCNT2* loci, but neither gene is expressed. (**c**) Hi-C interaction density mapped onto the ecDNA sequence. Long-range chromatin interactions spanning breakpoint junctions are indicated by arrows. (**d**) Gene expression in the MB268 primary tumor. All ecDNA-amplified genes are indicated by the orange swarmplot; highly expressed genes are labelled. The violin plot indicates a kernel density estimate of the distribution of expression of all genes in MB268. (**e**) FISH of *MDM4* in an FFPE slide of MB268 primary tumor confirms extrachromosomal amplification of *MDM4*.

